# Riboflow: using deep learning to classify riboswitches with ~99% accuracy

**DOI:** 10.1101/868695

**Authors:** Keshav Aditya R. Premkumar, Ramit Bharanikumar, Ashok Palaniappan

## Abstract

Riboswitches are cis-regulatory genetic elements that use an aptamer to control gene expression. Specificity to cognate ligand and diversity of such ligands have expanded the functional repetoire of riboswitches to mediate mounting apt responses to sudden metabolic demands and signal changes in environmental conditions. Given their critical role in microbial life, and novel uses in synthetic biology, riboswitch characterisation remains a challenging computational problem. Here we have addressed the issue with advanced deep learning frameworks, namely convolutional neural networks (CNN), and bidirectional recurrent neural networks (RNN) with Long Short-Term Memory (LSTM). Using a comprehensive dataset of 32 ligand classes and a stratified train-validate-test approach, we demonstrated the superior performance of both the deep models (CNN and RNN) relative to other conventional machine learning classifiers on all key performance metrics, including the ROC curve analysis. In particular, the bidirectional LSTM RNN emerged as the best-performing learning method for identifying the ligand-specificity of riboswitches with an accuracy > 0.99 and macro-averaged F-score of 0.96. A dynamic update functionality is inbuilt to account for the discovery of new riboswitches and extend the predictive modelling to any number of new additional classes. Our work would be valuable in the design and assembly of genetic circuits and the development of the next generation of antibiotics. The software is freely available as a Python package and standalone resource for wide use in genome annotation and biotechnology workflows.

**Availability:** PyPi package: riboflow @ https://pypi.org/project/riboflow

Repository with Standalone suite of tools: https://github.com/RiboswitchClassifier

Language: Python 3.6 with `numpy, keras`, and `tensorflow` libraries.

Licence: MIT

## Introduction

Riboswitches are ubiquitous and critical metabolite-sensing gene expression regulators in bacteria that are capable of folding into at least two alternative conformations of 5’UTR mRNA secondary structure, which functionally switch gene expression between on and off states [1-3]. The selection of conformation is dictated by the binding of ligand cognate to the aptamer domain of a given riboswitch [4-6]. Cognate ligands are key metabolites that mediate responses to metabolic or external stimuli. Consequent to conformational changes, riboswitches weaken transcriptional termination or occlude the ribosome binding site thereby inhibiting translation initiation of associated genes [7-8]. Riboswitches provide an intriguing window into the ‘RNA world’ biology [9-12] and there is evidence of their wider distribution in complex genomes [13-16]. The modular properties of riboswitches have engendered the possibility of synthetic control of gene expression [17], and combined with the ability to engineer binding to an *ad hoc* ligand, riboswitches have turned out to be a valuable addition to the synthetic biologist’s toolkit [18,19]. In addition to orthogonal gene control they are useful in a variety of applications, notably metabolic engineering [20], biosensor design [21,22] and genetic electronics [23]. Riboswitches have been used as basic computing units of a complex biocomputation network, where the concentration of the ligand of interest is titrated into measurable gene expression [24,25]. Riboswitches have also been directly used as posttranscriptional and translational checkpoints in genetic circuits[26]. Their key functional roles in infectious agents but absence in host genomes make them attractive targets for the design of cognate inhibitors [27-29]. Characterisation of riboswitches would lead to the rapid assemblies of reliable genetic circuits. In view of their myriad uses, a robust computational method for the accurate characterisation of novel riboswitch sequences would be of great interest.

Since the discovery of riboswitches [30,31], many computational efforts have been advanced for their characterisation, notably Infernal [32], Riboswitch finder [33], RibEx [34], RiboSW [35] and DRD [36], and reviewed in Clote [37] and Antunes et al [38]. These methods used probabilistic models of known classes with or without secondary structure information to infer or predict the riboswitch class. Singh and Singh explored featuring mono-nucleotide and di-nucleotide frequencies in a supervised machine learning framework to classify different riboswitch sequences, and concluded that the multi-layer perceptron was optimal [39]. Their work achieved modest performance (F-score of 0.35 on 16 different riboswitch classes). None of the above methods were shown to generalise well to unseen riboswitches. Our remedy was to explore the use of deep learning models for riboswitch classification. Deep networks are relatively recent neural network-based frameworks that are being used to great success in biomedical research. Convolutional neural networks are one type of deep learning, known for hierarchical information extraction. Such architectures with alternating convolutional and pooling layers have been earlier used to extract structural and functional information from genome sequences [40-43]. Recurrent neural networks are counterparts to CNNs and specialise in extracting features from time-series data [44]. RNNs with Long Short-Term Memory (termed LSTM) incorporate feedback connections to model long-term dependencies in sequential information [45], such as in speech and video [46]. This feature of LSTM RNNs immediately suggests their use in character-level modelling of biological sequence data [47,48]. Bidirectional LSTM RNN have been shown to be especially effective, given that they combine the outputs of two LSTM RNNs, one processing the sequence from left to right, the other one from right to left, together enabling the capture of dynamic temporal or spatial behavior [49]. Bidirectional LSTM RNNs are a particularly powerful abstraction for modelling nucleic acid sequences whose spatial secondary structure determines function [50]. Here we have evaluated the relative merits of a spectrum of state-of-the-art learning methods for resolving the ligand-specificity from riboswitch sequence.

## Methods

### Dataset and pre-processing

We searched the Rfam database of RNA families [51] with the term “Riboswitch AND family” and the corresponding hit sequences were obtained in fasta format from the Rfam ftp server (Rfam v13 accessed on July 6, 2019). Each riboswitch is represented by the coding strand sequence, with uracil replaced by thymine, thereby conforming to the nucleotide alphabet ‘ACGT’. Each sequence was scanned for nonstandard letters (i.e, other than the alphabet) and such occurrences were corrected using the character mapping defined in Table 1. The feature vectors for machine learning were extracted from the sequences. For each sequence, 20 features were computed, comprising four mononucleotide frequencies (A,C,G,T) and 16 dinucleotide frequencies. Deep models, namely convolutional neural networks (CNNs) and bidirectional recurrent neural networks with LSTM (hereafter simply referred as RNNs) are capable of using the sequences directly as the feature space, obviating any need for feature engineering. The data for each instance consists of the sequence, its 20 features and the riboswitch class. Python scripts used to create this final dataset are available at https://github.com/RiboswitchClassifier.

**Table 1.** Non-standard nucleotide mapping. Rare occurrences of non-standard nucleotides in the sequences were converted using this mapping key.

| S. No. | Original letter | Mapped character | #occurrences in dataset |
| --- | --- | --- | --- |
| 1 | R | G | 6 |
| 2 | Y | T | 8 |
| 3 | K | G | 1 |
| 4 | S | G | 3 |
| 5 | W | A | 2 |
| 6 | H | A | 2 |

### Predictive Modelling

The machine learning problem is simply stated as: given the riboswitch sequence, predict the ligand class of the riboswitch. Towards this, a battery of eight supervised machine learning and deep classifiers were studied and evaluated in the present work, based on their algorithmic complementarity (Table 2). Classifiers derived from implementations in the Python scikit-learn machine learning library [52] (www.sklearn.com) are referred to as base models and include the Decision Tree, K-nearest Neighbours, Gaussian Naive Bayes, the ensemble classifiers AdaBoost and Random Forest and the Multi-layer Perceptron. The deep classifiers namely CNN and RNN derived from implementations in the Python keras library (http://keras.io) on tensorflow [53]. For both the base and deep classifiers, the dataset was split into 0.9:0.1 training:test sets. Multi-class modelling is fraught with overfitting to particular classes (especially pronounced in cases of extreme class skew). To address this issue, the splitting process was stratified on the class, which ensured that each class was proportionately distributed in both the training and test sets.

**Table 2.** A summary of the riboswitch dataset used in our study. The dataset includes a mixture of metabolite/ion/cofactor/amino-acid/nucleotide/vitamin/signaling-molecule aptamer ligands. ‘Label no.’ corresponds to the response labels to be learnt in machine learning. The average length of all sequences in a given class is also given.

| Label no. | Rfam ID | Class Name | Class size | Avg. length |
| --- | --- | --- | --- | --- |
| 1 | RF00504 | Glycine riboswitch | 4592 | 100 |
| 2 | RF01786 | Cyclic di-GMP-II riboswitch | 661 | 86 |
| 3 | RF01750 | ZMP/ZTP riboswitch | 1674 | 92 |
| 4 | RF00059 | Thiamine pyrophosphate riboswitch | 12559 | 110 |
| 5 | RF01057 | SAH Riboswitch | 832 | 92 |
| 6 | RF01725 | SAM -1 -4 Variant riboswitch | 793 | 104 |
| 7 | RF00162 | SAM - 1 Riboswitch | 6027 | 113 |
| 8 | RF00174 | Cobalamin riboswitch | 14212 | 189 |
| 9 | RF01055 | Molybdenum Co-factor riboswitch | 1221 | 134 |
| 10 | RF01727 | SAM-SAH Riboswitch | 240 | 50 |
| 11 | RF01482 | AdoCbl riboswitch | 182 | 137 |
| 12 | RF03057 | nhaA-I RNA | 559 | 56 |
| 13 | RF01734 | Fluoride Riboswitch | 2018 | 70 |
| 14 | RF00167 | Purine Riboswitch | 2632 | 101 |
| 15 | RF00234 | Glucosamine-6-phosphate riboswitch | 936 | 175 |
| 16 | RF01739 | Glutamine riboswitch | 1103 | 64 |
| 17 | RF03072 | raiA RNA | 736 | 219 |
| 18 | RF03058 | sul1 RNA | 344 | 56 |
| 19 | RF00380 | Ykok riboswitch (Magnesium sensing) | 1059 | 170 |
| 20 | RF00168 | Lysine Riboswitch | 2240 | 180 |
| 21 | RF03071 | DUF1646 | 265 | 53 |
| 22 | RF01689 | AdoCbl variant RNA | 212 | 125 |
| 23 | RF00379 | ydaO/yuaA leader | 3918 | 164 |
| 24 | RF00634 | SAM - 4 Riboswitch | 1245 | 116 |
| 25 | RF01767 | SAM - 3 Riboswitch | 195 | 90 |
| 26 | RF00080 | yybP-ykoY manganese riboswitch | 833 | 158 |
| 27 | RF02683 | NiCo riboswitch (Nickel or Cobalt sensing) | 235 | 97 |
| 28 | RF00442 | Guanidine-I riboswitch | 902 | 109 |
| 29 | RF00522 | Pre-queosine riboswitch -1 | 533 | 45 |
| 30 | RF00050 | Flavin Mononucleotide Riboswitch | 4080 | 142 |
| 31 | RF01831 | THF riboswitch | 663 | 102 |
| 32 | RF00521 | SAM - 2 Riboswitch | 819 | 78 |

### Evaluation Metrics

The performance of each classifier on the test set was evaluated using the receiver operating characteristic (ROC) analysis in addition to standard metrics such as the precision, recall, accuracy and F-score (harmonic mean of precision and recall) [54]. The ROC curve was obtained by plotting the TP rate vs. the FP rate i.e, sensitivity vs (1 – specificity), and the area under the ROC curve (AUROC) could be estimated to rate the model’s performance. AUROC represents the probability that a given classifier would rank a randomly chosen positive instance higher than a randomly chosen negative one. ROC analysis is robust to class imbalance, typical of the machine learning problem at hand, however a multi-class adaptation of the binary ROC is necessary. For each classifier, this is achieved by computing classwise binary AUROC values in a one-vs-all manner, followed by aggregating the classwise AUROC values into a single multi-class AUROC measure [55,56]. Aggregation of the classwise AUROC values could be done in atleast two ways:

1. micro-average AUROC,where each *instance* is given equal weight.
2. macro-average AUROC, where each *class* is given equal weight.

Both the micro-average and macro-average AUROC measures were used to evaluate the relative performance of all our classifiers.

### Dynamic extension of the models

Genome sequencing of diverse, exotic prokayotes is likely to yield new regulatory surprises mediated by riboswitches [57]. A model that could classify a fixed set of 32 classes remains static in the wake of exponentially growing number of known genomes. To address the challenge of extending the model to new classes, we have formulated a model updation strategy. The dynamic extension of the model is initiated by feeding the sequences corresponding to the new class(es) to an updater script, which revises the dataset and then trains a new model. The automation of modelling would ensure a user-friendly pipeline for the generative modelling of any number of riboswitch classes along with the production of performance metrics of such models. Such an automation has been implemented to extend the eight classifiers compared to model any number of new riboswitch classes. The implementation includes a mechanism for hyperparameter optimization, relieving any demands on the user. The updater script and allied features are available at https://github.com/RiboswitchClassifier.

## Results

Our Rfam query retrieved 39 riboswitch class hits, however seven of these classes had a membership of less than 100 sequences and were filtered out. Subsequently, our dataset consisted of 32 riboswitch classes with a total of 68,520 sequences. A summary of this dataset is presented in Table 2. The largest classes include the cobalamin and thiamine pyrophosphate classes, with > 10,000 riboswitches in each, accounting for considerable diversity within classes. Classes with >4,000 members include Flavin mononucleotide (FMN) and glycine riboswitches. Other notable classes with at least 1,000 members include the lysine, purine, fluoride and glutamine switches. The riboswitch sequences were inspected for the standard alphabet (Table 1) and the final pre-processed comma-separated values (csv) datafile with each instance containing the sequence, 20 features and riboswitch class was prepared (available at https://github.com/RiboswitchClassifier in the Datasets folder).

Table 3 summarises the classifiers used in the study. We noticed poor performance of the base models on the test set with default model parameters, which could be traced to persistent overfitting (dominated by the larger classes), despite stratifying both train and test sets on class. Hyperparameter optimisation of the default parameters is one solution to address this problem and was carried out on both base and deep models. Hyperparameter finetuning for each classifier was achieved by exhaustive grid search on the range of choices for all hyperparamaters of interest for that classifier. The grid search was evaluated with 10-fold cross-validation of the training set. This yielded the optimal hyperparameters for each classifier. The scripts for hyperparameter optimisation of the base models are available at https://github.com/RiboswitchClassifier. In the case of the deep models, hyperparameter optimisation was inbuilt into keras/TensorFlow model-building by setting the ‘validation’ flag to 0.1 during the training phase. This is essentially equivalent to a 10-fold cross-validation procedure. The exercise is summarized in Supplementary File S1, which includes the best configuration of the hyperparameter space for all the base and deep classifiers.

**Table 3.** Description of the base model and deep classifiers. Hyperparameters noted for each classifier are meant to be representative. For the deep models, any long riboswitch genome sequences were truncated to 250 nucleotides, which is an adjustable parameter (max_len) much larger than the average length of any riboswitch class. The Python3 library used for implementation of machine learning model is noted.

| Classifier | Features used | Hyperparameters of interest | ML Library |
| --- | --- | --- | --- |
| Decision Tree | 1- and 2-mer frequencies | Maximum features, Minimum sample split, Minimum sample leaf, Random state, Max depth | Sklearn |
| Gaussian Naïve Bayes | 1- and 2-mer frequencies | Priors | Sklearn |
| k-Nearest Neighbours | 1- and 2-mer frequencies | Number of neighbours, Leaf size, Weights, Algorithm | Sklearn |
| Adaptive Boosting | 1- and 2-mer frequencies | Number of estimators, Learning rate, Algorithm | Sklearn |
| Random Forest | 1- and 2-mer frequencies | Number of estimators, Max depth, Impurity criterion | Sklearn |
| Multi-layer Perceptron | 1- and 2-mer frequencies | Activation, Solver, Alpha (regularisation term), Learning rate, epochs, hidden layers, nodes per hidden layer | Sklearn |
| CNN | Riboswitch sequence | Number of filters, Kernel size, Activation function, Pooling method, number of Conv1D layers, Dropout ratio, Optimiser, #Epochs | TensorFlow (Keras) |
| Bidirectional RNN with LSTM | Riboswitch sequence | Activation function, number of LSTM nodes, number of Bidirectional layers, Dropout ratio, Optimiser, #Epochs | TensorFlow (Keras) |

The optimised CNN and RNN architectures are illustrated in Figure 1. In the CNN, two convolutional layers were used followed by a pooling layer and dropout layer before flattening to a fully connected output layer. The RNN employed two sophisticated bidirectional LSTM units sandwiched by dropout layers before flattening to a fully connected output layer. The number of training epochs necessary for each deep model was determined based on the convergence of the error function (shown in Figure 2).

**Figure 1.**
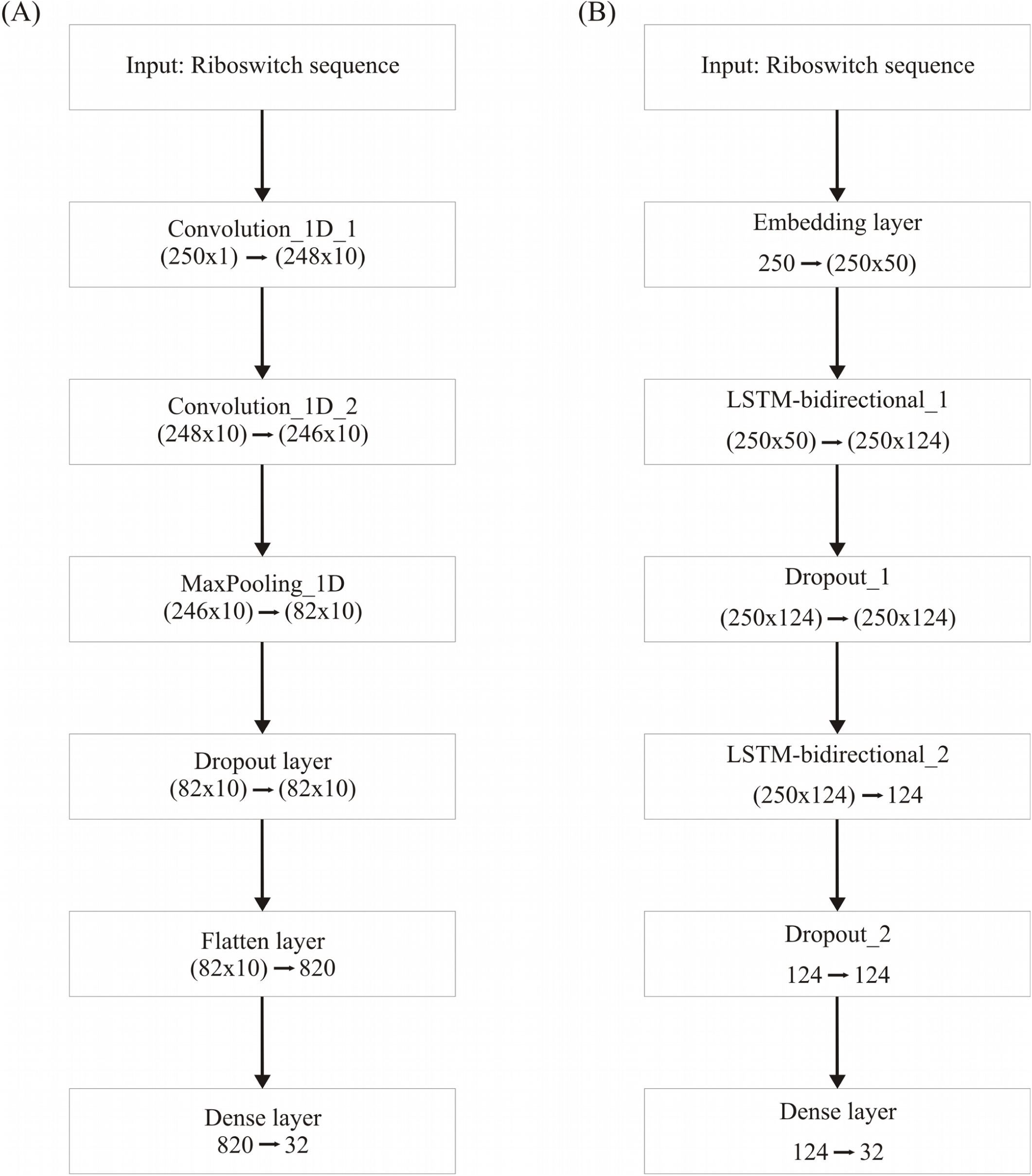
Deep learning frameworks used in the study. (A), CNN architecture, optimised for two 1-dimensional convolutional layers; and (B), Bidirectional RNN with LSTM, optimised for two bidirectional layers. Two dropout layers are used in the RNN

**Figure 2.**
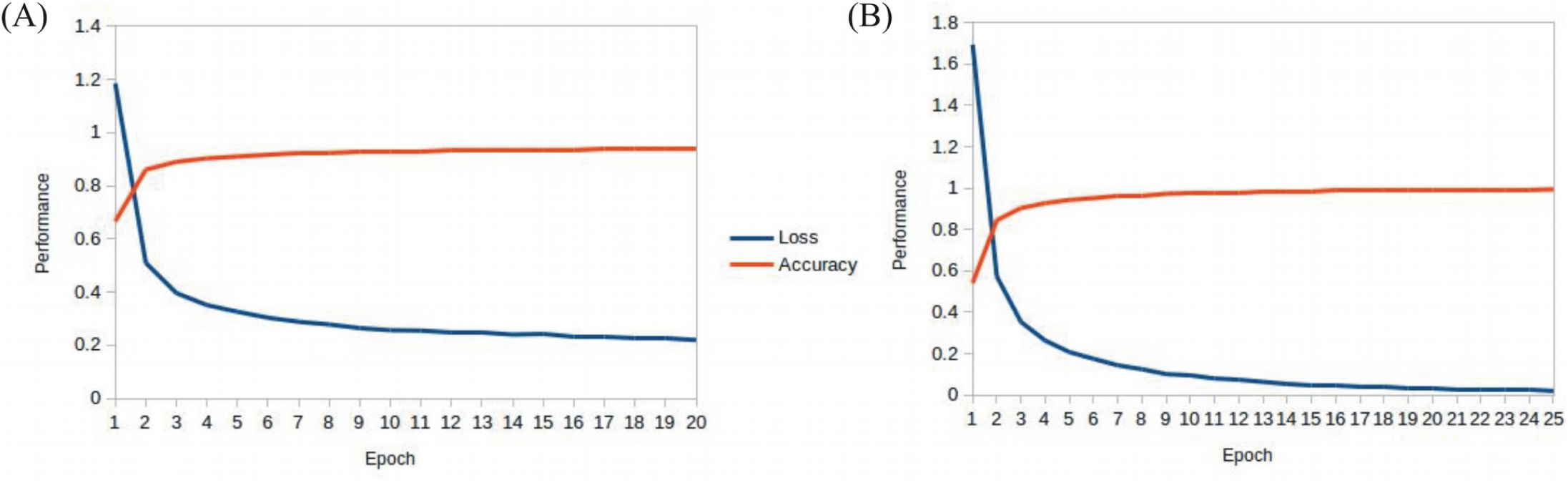
Epoch tuning curves for the CNN (A) and RNN (B). The CNN converges faster with respect to the number of epochs, however the RNN learns better, as seen with the decreasing loss function.

With the optimised classifiers, the performance of the predictive modelling was evaluated on the unseen testing set. Figure 3 shows the resultant classifier performance by an array of metrics including accuracy, and F-score. It is abundantly clear that the deep models vastly outperformed the base classifiers in all metrics across all classes. Figures 4, 5 show the ROC curves along with the micro-averaged and macro-averaged AUROC for the base models and the deep models, respectively. The AUROC is indicative of the quality of the overall model. It is seen that the AUROC is 1.00 for all classes for the RNN. Table 4 summarises the performance of the classifiers, with the detailed classwise F-score of each classifier available in the Supplementary Table S2 and the classwise break-up of the AUROC of all classifiers in the Supplementary Table S3.

**Table 4.** Performance metrics for all classifiers. The macro-averaged values of precision, recall and F-score are shown. Micro AUROC, micro-average AUROC; Macro AUROC, macro-average AUROC.

| Model | Accuracy | Precision | Recall | F-score | Micro AUROC | Macro AUROC |
| --- | --- | --- | --- | --- | --- | --- |
| Decision Tree | 0.54 | 0.49 | 0.39 | 0.42 | 0.88 | 0.81 |
| Gaussian NaïveBayes | 0.37 | 0.39 | 0.46 | 0.38 | 0.9 | 0.89 |
| K-neighbors | 0.67 | 0.75 | 0.55 | 0.61 | 0.94 | 0.88 |
| AdaBoost | 0.47 | 0.42 | 0.36 | 0.36 | 0.89 | 0.92 |
| Random Forest | 0.71 | 0.86 | 0.58 | 0.65 | 0.98 | 0.97 |
| Multi-layer perceptron | 0.74 | 0.75 | 0.67 | 0.70 | 0.99 | 0.98 |
| CNN | 0.97 | 0.98 | 0.91 | 0.93 | 1 | 1 |
| RNN | 0.99 | 0.97 | 0.96 | 0.96 | 1 | 1 |

**Figure 3.**
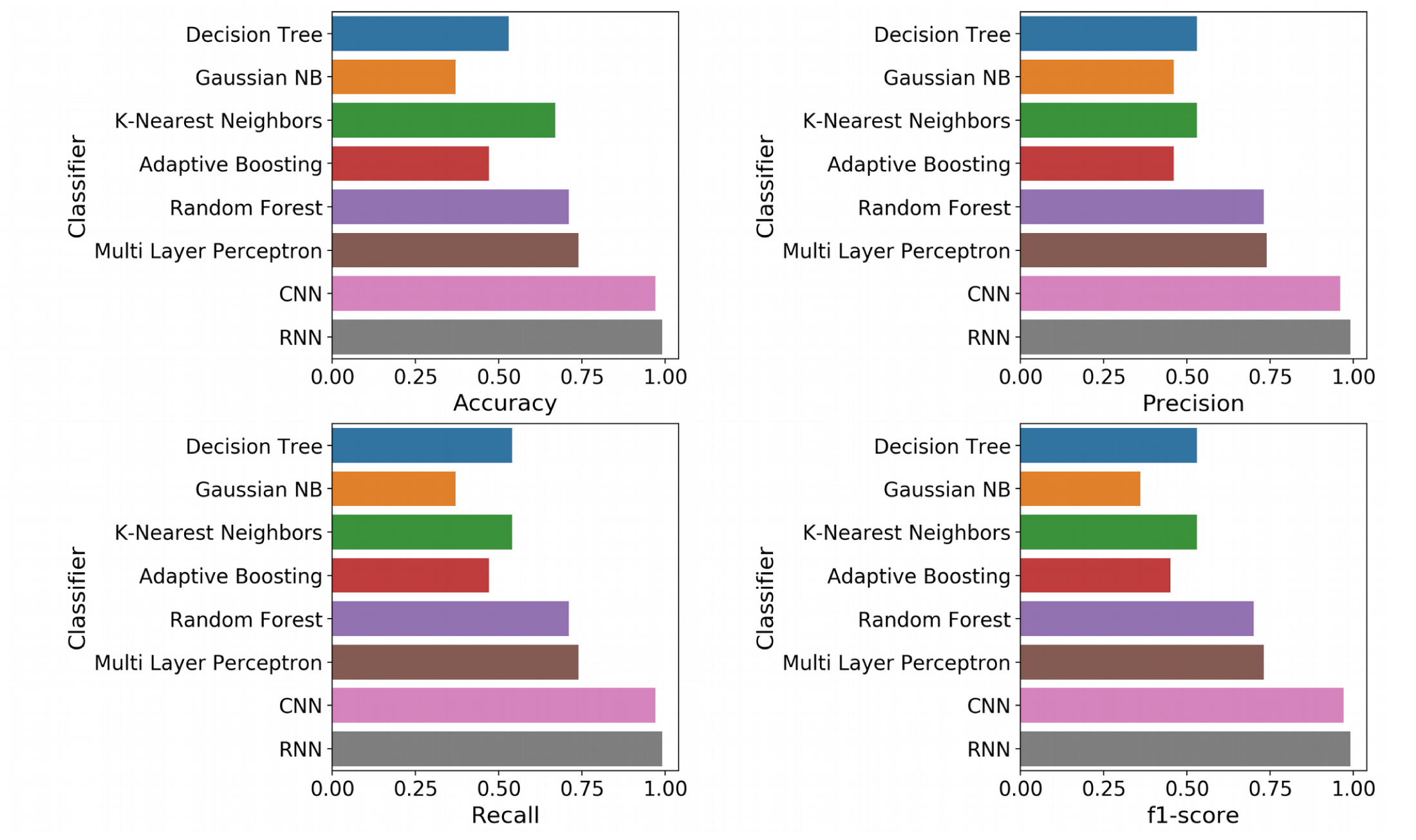
Standard performance metrics. Clockwise from top left, Accuracy; Precision; F-score; and Recall. The overall precision, recall and F-score were computed by macro-averaging the classwise scores. The deep models emerged as vastly superior alternatives to the base machine learning models on all performance metrics.

**Figure 4.**
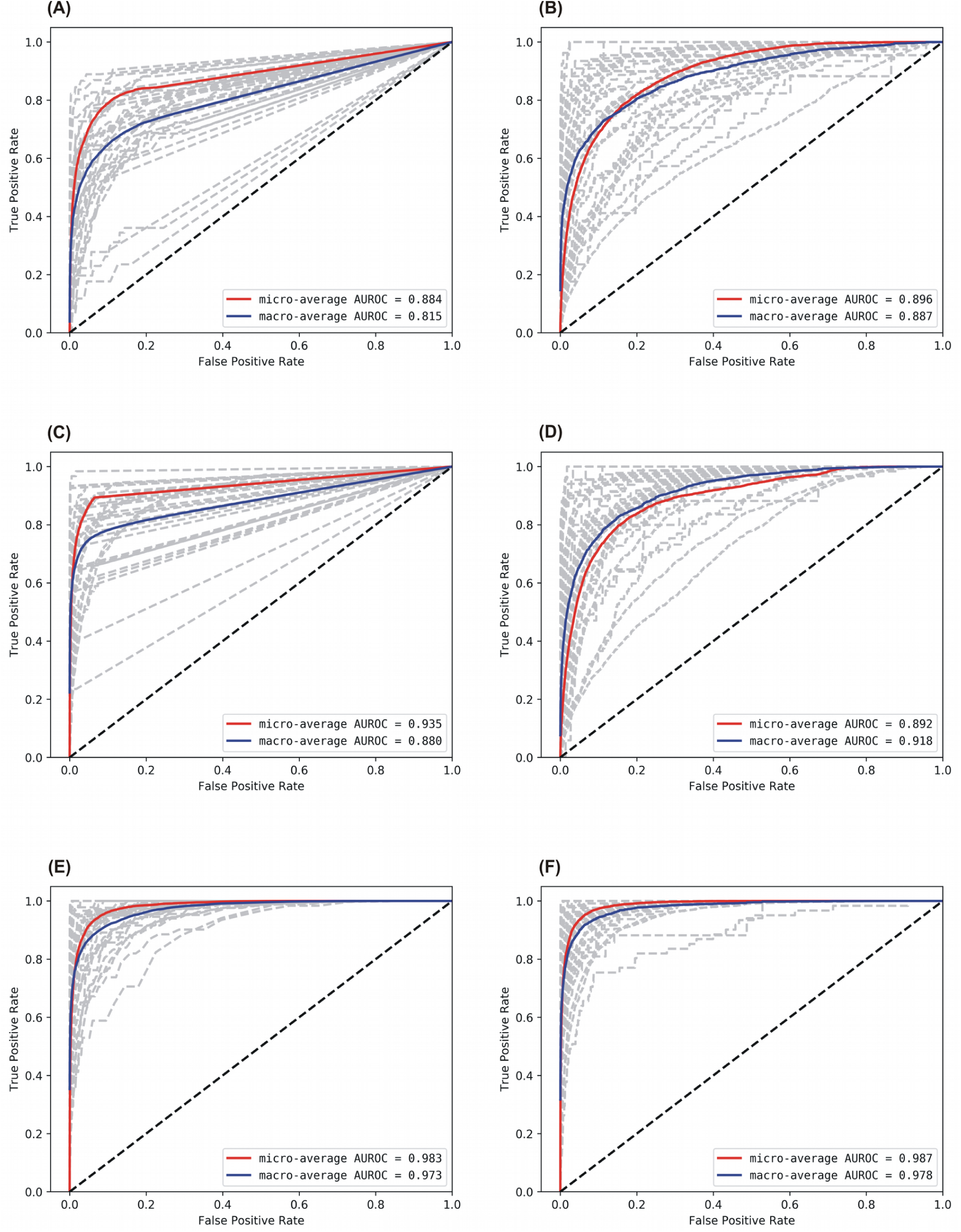
AUROC for the base models. A: Decision Tree, B: Gaussian NB, C: kNN, D: AdaBoost, E: Random Forest, and F: Multi-layer Perceptron. Grey lines denote classwise AUROCs of all 32 classes, from which it is clear that not all classes are equally learnt.

**Figure 5.**
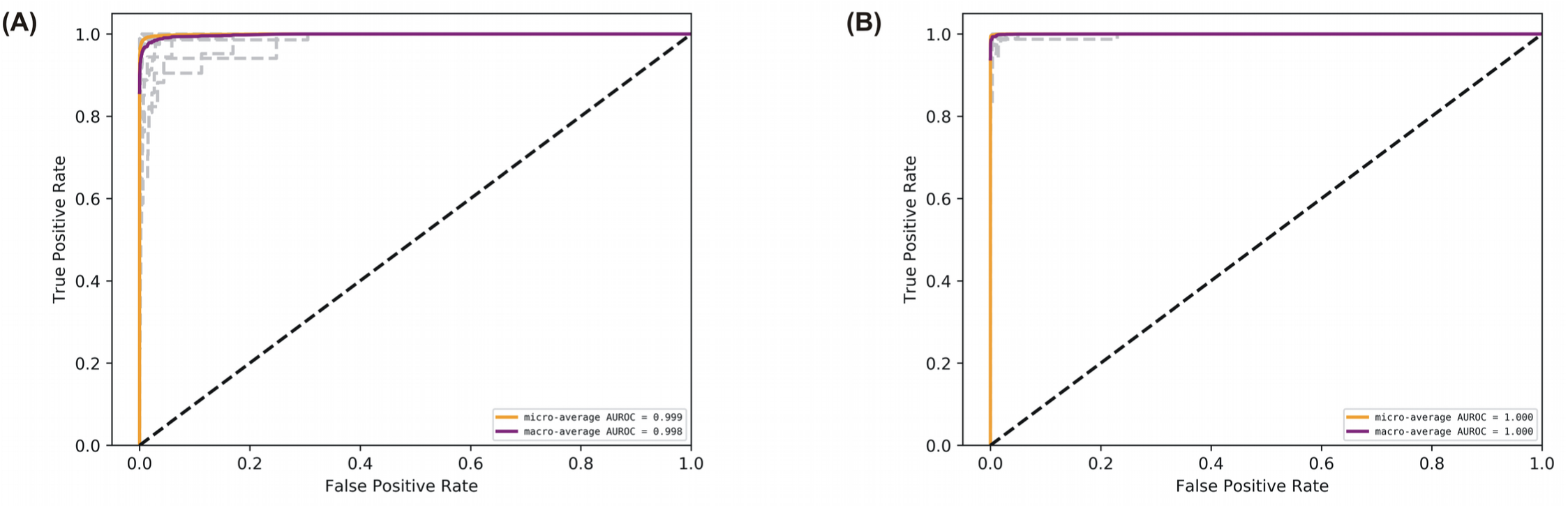
AUROC for the deep models. A: CNN, and B: RNN. Grey lines denote classwise AUROCs of all 32 classes. It clear that RNN achieves learning perfection at both the macro and classwise levels.

## Discussion

The RNN model marginally (but clearly) outperformed the CNN model, and both of them significantly outperformed all the base models on all key metrics, notably accuracy and F-score. The best-performing among the base models was the Multi-layer Perceptron. It is noteworthy that the AUROC approached unity and near-perfection for both the deep models, especially the bidirectional RNN with LSTM. This implied that the use of k-mer features masked long-range information whereas the deep models were able to capture such correlations directly from the full sequence. These results affirmed that RNNs could be used to effectively simulate the interactions in biological sequences.

The F-score (a measure of balanced accuracy) is a more unforgiving metric than AUROC in the case of multi-class problems(Table 4). While the CNN and RNN had macro-averaged F-scores of 0.93 and 0.96 respectively, none of the base models exceeded 0.70 including the multilayer perceptron. Supplementary Table S2 provides the classwise F-scores of all classifiers and afforded acute analysis. All the base models struggled to classify the largest riboswitch classes, namely TPP and Cobalamin. This is a consequence of the diversity of such large riboswitch classes, making the ‘outlier’ members harder to classify correctly. These diverse classes did not pose any problems for the deep models. The greatest challenges to both base and deep models were the AdoCbl and AdoCbl variant riboswitch classes, but even here the RNN showed remarkable consistency relative to all other classifiers. The other classes that were significantly challenging to the base models but not to the deep models included Cyclic di-GMP-II, ZMP/ZTP, SAM 1-4 variant, Molybdenum co-factor, Glucosamine-6-phosphate and Guanidine-I riboswitch classes. It is seen that the classification problems arise at the extremes of class sizes. Too large the class, the diversity is challenging, whereas too small and the learning itself is incomplete and challenged. The deep models – RNN and CNN – consistently performed well across all classes, independent of the size of the class. It could be inferred in this case that using direct features (i.e, sequences) rather than engineered features (i.e, k-mer frequencies) led to more robust models.

These results might be put in perspective by benchmarking against the existing literature. Guillen-Ramirez and Martinez-Perez [58] extended the k-mer features logic and arrived at an optimal combination of 5460 k-mer features. Using a limited dataset of 16 classes, they used state-of-the-art machine learning to achieve accuracies in the high nineties, however their results did not generalise equally to riboswitches with remote homology. For e.g, their best-performing classifier (Multi-layer Perceptron) misclassified 6 out of the 225 instances of Lysine riboswitch as cobalamin-gated. The source code for the features and modelling used in their work is not available for reproducible research.

An interesting benchmark is afforded by the Riboswitch Scanner [59], which used profile HMMs [60] of riboswitch classes to detect riboswitches in genomic sequences. While our method addresses inter-class discrimination of riboswitch sequences, Riboswitch Scanner is a web-server that essentially performs riboswitch vs not-riboswitch discrimination for user-given riboswitch classes. The absence of F-score metrics does not allow for direct comparisons, however the sensitivity and specificity seemed consistently comparable for most classes, except the Glycine, THF and SAM I-IV variant riboswitch classes. It must be noted that their method is validated with Rfam seed sequences, without consideration for the proliferation of riboswitch sequences. Performance evaluation on limited data could inflate performance estimates and complicate their interpretation.

It must be noted that riboswitches are precisely specific to cognate ligands. For e.g, the AdoCbl riboswitch would not tolerate a methyl-substituted cobalamin [61] nor does the TPP riboswitch interact with thiamine or thiamine monophosphate [62]. At the same time, these two riboswitches are very diverse in their phylogenomic distribution and actual sequences. The key to effective learning lies in treading a fine balance between the intra-class diversity and inter-class specificity. It is remarkable the bidirectional LSTM RNN was able to achieve exactly this tradeoff. The roots of such performance of the deep models in general has been recently speculated to be related to the lottery ticket hypothesis [63] as well as learning the intrinsic dimension of the problem [64], here the classification of riboswitches.

To extend the functionality of our work, we have introduced a dynamic component to all our models, both base and deep. With the exponential growth in genome sequencing, the room for riboswitch discovery is enormous. Our models could accommodate new riboswitch class definitions by way of dataset augmentation, thereby making them general and more robust. This work used the said dynamic functionality to extend a preliminary 24-class model to the present 32 classes with not the slightest impact on the performance. The dynamic functionality is provided by the script dynamic.py available at https://github.com/RiboswitchClassifier. The script has two use cases depending on the number of new classes.

~~~
#Use case 1.
$ dynamic.py -fa riboclass.fa
#Use case 2.
$ dynamic.py -d dir
~~~

In the simpler use case, a single new class is added to the dataset by specifying the option ‘-fa’ and providing the name of the fasta datafile. Alternatively, multiple new class definitions could be added to the model by specifying the ‘-d’ option and providing the name of the directory that contains one fasta datafile for each new class. The script processes the datafiles for compatibility and the alphabet, and adds them to the existing dataset. The updated dataset is read as the input for training the new base models, and the CNN and RNN deep models. The new models would then be able to handle any number of new riboswitch classes, regardless of sequence diversity considerations. Given that the performance of the deep models remained unaffected by an increase in the number of classes for learning, the models could be expected remain robust as even more riboswitch classes are discovered.

Training the CNN on an Intel i7 processor @3.4GHz with 8GB RAM took ∼ 8 minutes, whereas training the RNN took about 5 hours. In the event of many new riboswitch classes, our recommendation would be to dynamically update the CNN model (`cnnApp.py`) available at https://github.com/RiboswitchClassifier. The trained deep models are available in hdf5 format at https://github.com/RiboswitchClassifier, and are modest in size (CNN model: 233 Kb; RNN model: 1.8 Mb). To make the workflow fully automatic, we have developed a Python package `riboflow` (https://pypi.org/project/riboflow/) mirroring the best RNN model. `riboflow` could be installed using the Python package installer, `pip (or pip3)`, and the dependencies `numpy, tensorflow` and `keras`. The following interactive code gives an example of `riboflow` uses.

~~~
>import riboflow
#Construct a Python list of riboswitch sequences. A sequence is a string in
alphabet ‘ATGC’
> sequences = [
“TTTTTTTTGCAGGGGTGGCTTTAGGGCCTGAGAAGATACCCATTGAACCTGACCTGGCTAAAACCAGGGTAGGGAATTGC”,
“CTCTTATCCAGAGCGGTAGAGGGACTGGCCCTTTGAAGCCCAGCAACCTACACTTTTTGTTGTAAGGTGCTAACCTGAGC”,
“CCACGATAAAGGTAAACCCTGAGTGATCAGGGGGCGCAAAGTGTAGGATCTCAGCTCAAGTCATCTCCAGATAAGAAATA”
]
#Use case 1. Return the most probable class for each riboswitch sequence:
> riboflow.predict(sequences, “predict_class”)
#Use case 2. Return the complete vector of class probabilities for each riboswitch
sequence, to disambiguate potential class confusion:
> riboflow.predict(sequences, “predict_prob”)
~~~

In summary, we have developed `riboflow`, a python package as well as standalone suite of tools, that have been validated and thoroughly tested on 32 riboswitch classes. By using large and complete datasets, the variance of our modelling procedure has been optimised and this ensures the generality and applicability of our models on new instances without compromise of performance. `riboflow` is an off-the-shelf solution that would afford the ready programmatic incorporation of the RNN model into automatic annotation pipelines. Should the user wish to use any of our other trained models, `pickled` models (https://docs.python.org/3/library/pickle.html) are available at https://github.com/RiboswitchClassifier. Our work presents an intuitive general-purpose platform for the effortless characterization of new riboswitch sequences, which presents applications in genome annotation and synthetic biology, including the rapid design of novel genetic circuits with exquisite specificity.

## Conclusion

We have demonstrated that CNN and RNN are capable of robust multi-class learning of ligand specificity from riboswitch sequence, with the RNN boasting an F-score of ∼0.96. The confidence of classification could be obtained from an inspection of the predicted classwise probabilities. The bidirectional LSTM RNN model has been packaged into `riboflow` to enable embedding into automated genome annotation pipelines and biotechnology workflows. The CNN shows the best tradeoff between the time-cost of training the model and overall performance and could be used for the task of learning new riboswitch classes using a dynamic update option that is provided. All the code used in our study is made freely available for use by the scientific community as well as in the interest of reproducible research. Our study has highlighted the use of macro-averaged F-score as a disciminating objective metric of classifier performance on multi-class data. Our work reaffirms the competitive advantages of bidirectional LSTM RNNs over conventional machine learning and hidden markov profiles in modelling data sequences, and opens up their applications for modelling other noncoding RNA elements. Riboswitches are novel and exciting targets for the development of new class of antibiotics and our work would help towards the design of riboswitch inhibitors to combat multi-drug resistant pathogens.

## Acknowledgements

This work was funded in part by DST-SERB grant EMR/2017/000470 and the SASTRA TRR grant (to A.P.). We are grateful to the School of Chemical and BioTechnology, SASTRA Deemed University for infrastructure and computing support. We would like to thank Nachal for help with figures.

## Author contributions

Conceived and designed the work: A.P. Performed experiments: KP, RB, AP. Analyzed data: AP, RB, KP. Wrote the paper: A.P.

