## Supplementary for "Riboflow: using deep learning to classify riboswitches with ~99% accuracy": File S1.pdf

### Hyperparameter tuning

**Supervised machine learning**
Grid search was performed in a non-randomized manner with different choices on the range of different hyperparameters specific to each model. 10-fold cross validation was used to estimate the best hyperparameters. Hyperparameter tuning wasn't performed on the Gaussian Naive Bayes model because no priors were specified to the probabilistic model.

| Model Name | Hyperparameters and Range | Accuracy before grid search(Test set) | Optimal Hyperparameter | Accuracy after Grid search(Test set) |
|---|---|---|---|---|
| Random Forest | Number of estimators:1000, 2000,3000,4000,5000<br><br>Max Depth:60,70,80, 90,100<br><br>Criterion : Gini,Entropy | 0.63 | Number of estimators: 3000<br><br>Max depth: 80<br><br>Criterion: Gini | 0.70 |
| Decision Tree | Maximum features: Auto, Sqrt, log2,None<br><br>Minimum sample split: 2,3,4,5,6,7,8,9,10,11,12,13,14,15<br><br>Minimum Sample Leaf:1,2,3,4,5,6,7,8,9,10,11<br><br>Random state:123,345,None<br><br>Max depth:5,10,15,20,25,None | 0.48 | Max Depth: 15<br><br>Max Features: None<br><br>Minimum sample leaf: 8<br><br>Minimum sample split: 3<br><br>Random state: None | 0.53 |

| | | | | |
|---|---|---|---|---|
| KNN | Number of neighbours: 5,6,7,8,9,10<br>Leaf size: 1,2,3,5<br>Weights: uniform, distance<br>Algorithm: auto, ball_tree,kd_tree,brute | 0.60 | Number of neighbours: 8<br>Leaf size: 1<br><br>Weights: Distance<br><br>Algorithm:Auto | 0.65 |
| AdaBoost | Number of Estimators: 1,50,1000,2000,3000<br><br>Learning Rate: 0.01,0.1,1.0,5.0,10.0<br><br>Algorithm: SAMME,SAMME.R | 0.24 | Number of Estimators: 1000<br><br>Learning Rate: 1.0<br><br>Algorithm: SAMME | 0.44 |
| Multi Layer Perceptron | Activation: tanh,relu<br><br>Solver: sgd,adam<br><br>Alpha:0.0001,0.01,0.1,0.05,1.0<br><br>Learning rate: constant,adaptive | 0.68 | Activation: Relu<br><br>Solver: Adam<br><br>Alpha :0.01<br><br>Learning rate: adaptive | 0.72 |

**Deep Learning:**
MLP Default: single hidden layer with 100 nodes (default).
Increasing the epoch, hidden layers, number of nodes per layer, reducing learning rate and changing the activation function did not significantly increase the accuracy and F1 score.
This led us to further explore other neural network architectures like the Recurrent Neural Network and Convoluted Neural Network.

Batch size for both: 128.
CNN:
Objective function: accuracy (scorer)
Gradually increased / changed the following
1. Number of filters, kernal size
2. Changing the activation function
2. Use of Average Pooling and MaxPooling
4. Increasing the number of Conv1D layers
5. Changing Dropout Ratio
6. Changing the Optimiser in use
7. Increasing Epochs

Saturation at 97 - 98% Test Accuracy was found when using 0.1 and 0.5 k fold cross validation split (Run Time in order of minutes < 30 minutes):
Filters = 10
kernal size = 3
Conv1D layers = 2
Epochs = 20
Optimiser = rmsprops
Activation = Relu
maxPooling
Dropout = 0.5

To achieve higher accuracy using the RNN :

Gradually increased / changed the following
1. Changing the activation function
2. Increasing number of LSTM nodes
3. Increasing the number of Bidirectional Layers
4. Changing Dropout Ratio
5. Changing the Optimiser in use
6. Increasing Epochs

Saturation at 99% Test Accuracy was found when using 0.1 and 0.5 k fold cross validation split (Run Time in order of Hours):

Bidirectional layers = 2
Epochs = 25
LSTM nodes = 62
Optimiser =adams
Activation = Relu
Dropout = 0.2
